# Slow running benefits: Boosts in mood and facilitation of prefrontal cognition even at very light intensity

**DOI:** 10.1101/2024.01.29.575971

**Authors:** Chorphaka Damrongthai, Ryuta Kuwamizu, Yudai Yamazaki, Naoki Aoike, Dongmin Lee, Kyeongho Byun, Ferenc Torma, Worachat Churdchomjan, Michal A. Yassa, Kazutaka Adachi, Hideaki Soya

## Abstract

Although running upright has been reported to have positive effects on both physical and mental health, the minimum running intensity/speed that would benefit mood and prefrontal cognition is not yet clear. For this reason, we aimed to investigate the acute effect of very slow running, which is classified as a very light intensity exercise, on mood, executive function (EF), and their neural substrates in the prefrontal cortex (PFC). Twenty-four healthy participants completed a 10-minute very slow running session on a treadmill at 35% 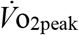 and a resting control session in randomized order. EF was measured using the Stroop task and the mood state was measured using the Two-Dimensional Mood Scale (TDMS) before and after both sessions. Cortical hemodynamic changes while performing the task were monitored using functional near-infrared spectroscopy (fNIRS). The results show that ten minutes of very slow running significantly enhanced mood, reduced Stroop interference time (i.e., enhanced EF), and elicited left lateral PFC activation. Moreover, head acceleration, the magnitude of up-and-down oscillations, was measured during running, and a significant positive correlation with pleasant mood was found. Head acceleration is a remarkable characteristic of running and may be one of the factors related to a pleasant mood induced by very slow running. In conclusion, the current study reveals that a single bout of running, even at very slow speed, elicits a pleasant mood and improved executive function with enhancing activation in prefrontal subregions. This shed light on the slow running benefits to brain health.

## INTRODUCTION

Running has an extremely long history as a means of locomotion for humans. Human beings can navigate long distances through bipedal running (Bramble & Lieberman, 2004). It is interesting to hypothesize that human upright bipedal running is also strongly tied to the evolutionary expansion of the Homo sapiens’ frontal lobes and higher cognitive functions (Schulkin, 2016). While, in modern times, physical inactivity is a common problem in most developed countries, running has been gaining popularity since the 70s and has been proposed as the most popular democratic, which is to say universally accessible, sport in the world. Its connection to human evolution and its rising popularity lead us to believe that running is a fundamental and engaging physical activity with high adherence.

Running decreases not only the risk of cardiovascular diseases in general but also has positive effects on mental and brain health. Our previous neuroimaging study showed that moderate-intensity running exercise (50% 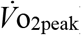) activated lateral prefrontal cortex (PFC) of both hemisphere and enhances executive function (Damrongthai *et al*., 2021). This resembles results from previous pedaling studies (Yanagisawa *et al*., 2010; Byun *et al*., 2014*b*; Kujach *et al*., 2018). Furthermore, Damrongthai *et al*. demonstrated a notable induction of a pleasant mood, although in previous cycling studies no such effect was found (Yanagisawa *et al*., 2010; Byun *et al*., 2014*b*; Kujach *et al*., 2018; Yamazaki *et al*., 2018; Damrongthai *et al*., 2021; Suwabe *et al*., 2021). However, moderate-intensity exercise is the intensity at which the hypothalamic stress response occurs (Soya *et al*., 2007). As such, vulnerable individuals, such as those with low fitness levels or the elderly, may find it difficult to adhere to a moderate-intensity exercise routine since moderate running is quite fast and generates a higher load on joints, which may lead to injury, compare to pedaling exercise. It is an intriguing challenge to see if a comparable effect can be achieved at a stress-free lower intensity, near walking speed. Our animal-to-human studies have shown that light-intensity exercise below the lactate threshold enhances the function of brain regions involved in cognition. In previous studies, pedaling exercise activated the left lateral PFC and improved executive function without significant changes in mood (Byun *et al*., 2014*b*; Kuwamizu *et al*., 2023). Based on this study, it is expected that even very slow running can improve executive function. Furthermore, if mood improves even with very light-intensity running (i.e., very slow running) as it does with moderate-intensity running, we can propose that very slow running is a beneficial exercise mode that has a positive effect on both mood and executive function with minimal effort. Although very slow running may enhance executive function while also improving mood, its exact beneficial effects on mood and executive function, and the specific brain loci that activate to provide a neural basis remain to be determined.

Running strengthens cardiopulmonary function, muscles, and bones by supporting one’s body weight alternately on each foot in a regular rhythm (Garofolini & Taylor, 2019). The mechanical impact of each rhythmic foot strike on the muscles of the entire body, especially the legs, has been shown to lead to physiological change (e.g., improved peripheral and central blood circulation) (Palatini *et al*., 1989; Lyngeraa *et al*., 2013). One rodent study has demonstrated that mechanical stress from vertical head acceleration (HA; the rate of vertical change in head velocity) during running induces the internalization of serotonin receptors in the PFC and corresponding behavioral changes (Ryu *et al*., 2020). Mechanical stress from vertical HA, caused by adequate rhythmic up-and-down oscillations, is a characteristic of running not present in pedaling. Walking upright while always having at least one foot on the ground at all times is expected to produce a similar but partial effect; however, the up and down motion resulting from each foot striking the ground with a large impact force caused by body weight is unique to running. These running characteristics may have a connection with running-specific beneficial effects on mood and PFC function.

Activation in the PFC can be measured using functional near-infrared spectroscopy (fNIRS), a non-invasive brain imaging instrument. fNIRS is compact enough that it can be used in real-life environments, such as a gym, or in a laboratory setting next to, for example, a large treadmill, allowing the measurement of activated brain regions even during acute running. In the last two decades, fNIRS has contributed to research on brain activity during locomotion, including walking (Miyai *et al*., 2001; Vitorio *et al*., 2017). Importantly, however, methodological difficulties must be considered: motion artifacts and skin blood flow may contaminate the fNIRS signals measured during locomotion (Miyazawa *et al*., 2013; Vitorio *et al*., 2017), making it challenging to measure pure cortical activity based on neuro-vascular coupling. To solve this problem, one of our previous studies focused on the enhancement of executive function post-treadmill-running (Damrongthai *et al*., 2021). We developed an experimental protocol to minimize non-cortical fNIRS signals by examining in detail the decay of relevant indicators after exercise (Yanagisawa *et al*., 2010; Byun *et al*., 2014*b*, 2014*a*; Kujach *et al*., 2018; Damrongthai *et al*., 2021). We have been testing how the prefrontal sub-region that coincides with the Color-word Stroop test (CWST), a representative task of inhibitory control, is affected by exercise using the virtual registration method (Tsuzuki *et al*., 2007; Yanagisawa *et al*., 2010; Tsuzuki & Dan, 2014; Damrongthai *et al*., 2021). Thus, here we hypothesize that this approach will enable us to elucidate the neural basis of how very slow running enhances executive function.

This study aims to elucidate the effects of acute very slow running on mood and executive function, and on prefrontal sub-region activation as a neural basis. In addition, we will quantify HA as a candidate causal factor for the beneficial effects of very slow running.

## METHODS

### Participants

Participants included 24 young adults who were Japanese native speakers. All of them were nonsmokers, were right-hand dominant, had normal or corrected-to-normal vision, and had normal color vision. No participant reported a history of neurological or psychiatric disorders or had any conditions requiring medical care. One additional participant was tested but was excluded from the final analyses because her behavioral data change (Stroop interference reaction-time difference during her participation in the running condition) fell outside the range of 3 standard deviations from the mean. The demographic data of the participants is shown in Table 1. The study was approved by the Institutional Ethics Committee of the University of Tsukuba (approval number: tai020-120) where the protocol corresponded to the latest version of the Helsinki Declaration. Written informed consent was obtained from all participants. This sample size was confirmed to be acceptable: post-hoc sensitivity analysis, computed using G*Power 3.1, was performed based on the current sample size (24 participants) and resulted in an alpha of 0.05 and power of 80%, demonstrating sufficient sensitivity for detecting *t*-test differences exceeding *d* = 0.60 (with a two-tailed alpha), which is consistent with our previous studies (Yanagisawa *et al*., 2010; Byun *et al*., 2014*b*; Damrongthai *et al*., 2021).

**Table 1.** Participant demographics data (n = 24).

|  | Male (n = 13) | Female (n = 11) |
| --- | --- | --- |
| Age (yr) | 23.38 ± 1.76 | 20.55 ± 2.07 |
| Weight (kg) | 65.31 ± 11.65 | 49.12 ± 4.05 |
| Height (cm) | 172.15 ± 5.34 | 159.06 ± 4.03 |
| BMI (kg/m<sup>2</sup>) | 22.03 ± 3.92 | 19.40 ± 1.39 |
| Graded exercise test |  |  |
| $\dot{V}O_{2peak}$ (mL/kg/min) | 48.24 ± 6.25 | 40.03 ± 3.84 |
| HR<sub>max</sub> (bpm) | 191.15 ± 8.43 | 190.00 ± 7.25 |
| R | 1.14 ± 0.03 | 1.13 ± 0.06 |
| RPE | 19.23 ± 0.60 | 19.27 ± 1.01 |
| Final treadmill speed (km/h) | 16.54 ± 2.40 | 13.55 ± 1.97 |

Presented values are mean ± SD.
BMI = Body Mass Index; $\dot{V}O_{2peak}$ = Peak oxygen uptake; HR<sub>max</sub> = Maximal Heart Rate; R = Respiratory exchange ratio; RPE = Rate of Perceived Exertion

### Procedure for main experiment

The main experiment had three phases. First, the maximal oxygen uptake (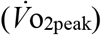) of each participant was measured to appropriately personalize the very light exercise intensity, which is to say treadmill speed, for very slow running. The target was 35% 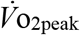 in accordance with the American College of Sports Medicine (Pescatello *et al*., 2014). Second, the proper timing for reassessing fNIRS measurements was determined since very slow running might induce non-cortical physiological signals, such as skin blood flow, that could consequently interfere with the fNIRS signal. It was determined that 5 min after very slow running was appropriate because within that time all physiological variables completely returned to baseline. Details of the first and second phases are provided in Supplemental Material 1 and Supplemental Material 2, respectively. Third, the effects of very slow running on mood, executive function, neural substrate, and HA were determined. All participants took part in two sessions, control (CON) and very slow running (SRUN), on separate days. The sessions were randomized and conducted with a counterbalance measure design. During the SRUN session, prefrontal hemodynamic changes during the CWST were measured using fNIRS before and 5 min after very slow running to avoid non-cortical physiological signal contamination. Participants were asked to slowly run on a treadmill at a personalized speed for 10 min. HA during running was assessed later, and the details are included in in Supplemental Material 3. Mood state was assessed before and after very slow running. In the CON session, participants were asked to sit on a chair on the treadmill instead of running for 10 min (Fig. 1). In addition, pupillometry was performed as an exploratory measurement item using a glass-type eye tracker, and the details can be found in Supplemental Material 4. It should be noted, however, that pupil measurements were limited to a small sample size due to device errors related to body movement.

**Figure 1.**
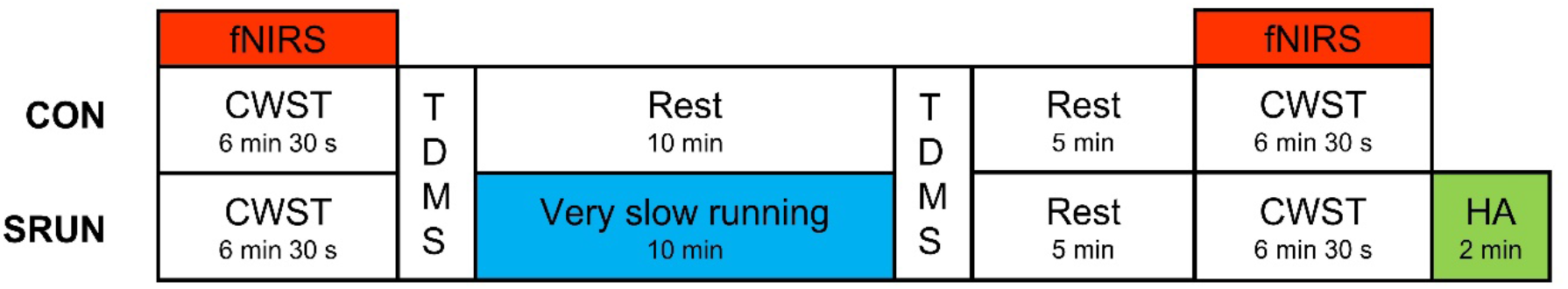
Experimental design for investigating the acute effect of very slow running on prefrontal cortical executive function. CON = control session, SRUN = very slow running session, CWST = color-word Stroop task, TDMS = Two-Dimensional Mood Scale, fNIRS = Functional near-infrared spectroscopy, HA = Head acceleration

### Mood measurements

The Two-Dimensional Mood Scale (TDMS) was adopted to evaluate mood state before and after both SRUN and CON conditions. The TDMS is a momentary mood scale consisting of eight words describing arousal and pleasure states: energetic, lively, lethargic, listless, relaxed, calm, irritated, and nervous. The participants used a 6-point Likert scale ranging from 0 = not at all to 5 = extremely to describe their feelings. Subsequently, the scores were calculated and interpreted to determine arousal and pleasure levels.

### Behavioral measurements

An event-related version of the CWST was used to evaluate executive function. The test was displayed on a monitor with two rows of letters. The participants were instructed to decide whether the color of the letters in upper row corresponded to the color name in the lower row. They were also asked to place their index fingers on “yes” and “no” buttons and to respond to the test by pressing the correct button as quickly as they could. Subsequently, reaction time (RT) and error rate (ER) were calculated. The CWST consisted of three conditions: neutral, congruent, and incongruent. For the neutral condition, the upper row displayed a row of X’s(XXXX) printed in red, green, blue, or yellow, and the lower row displayed the word ‘RED’, ‘GREEN’, ‘BLUE’, or ‘YELLOW’ printed in black. For the congruent condition, the upper row displayed the word ‘RED’, ‘GREEN’, ‘BLUE’, or ‘YELLOW’ printed in the congruent color (e.g., ‘BLUE’ was printed in blue), and the lower row displayed the same words as in the lower row of the neutral condition. For the incongruent condition, the upper row displayed a color word printed in an incongruent color to produce interference between the color word and the color name (e.g., ‘YELLOW’ was printed in red), and the lower row displayed the same words as in the lower row of the neutral and congruent conditions (Fig. 2A). Each experimental session consisted of 30 trials, made up of 10 neutral, 10 congruent, and 10 incongruent trials, which appeared in random order. The upper row was presented 100 ms before the lower row in order to shift visual attention. Each trial remained on the screen until the participant responded or for 2 s, whichever was shorter. Then, a fixation cross appeared on the screen as an inter-stimulus interval for 10-12 s to avoid prediction of the timing of the subsequent trial. Stroop interference, an index of executive function in the PFC, was calculated as the difference in reaction time between the incongruent and neutral conditions. All words were written in Japanese. The participants performed three practice sessions to ensure that they understood and were familiarized with the CWST well before starting the experiment.

**Figure 2.**
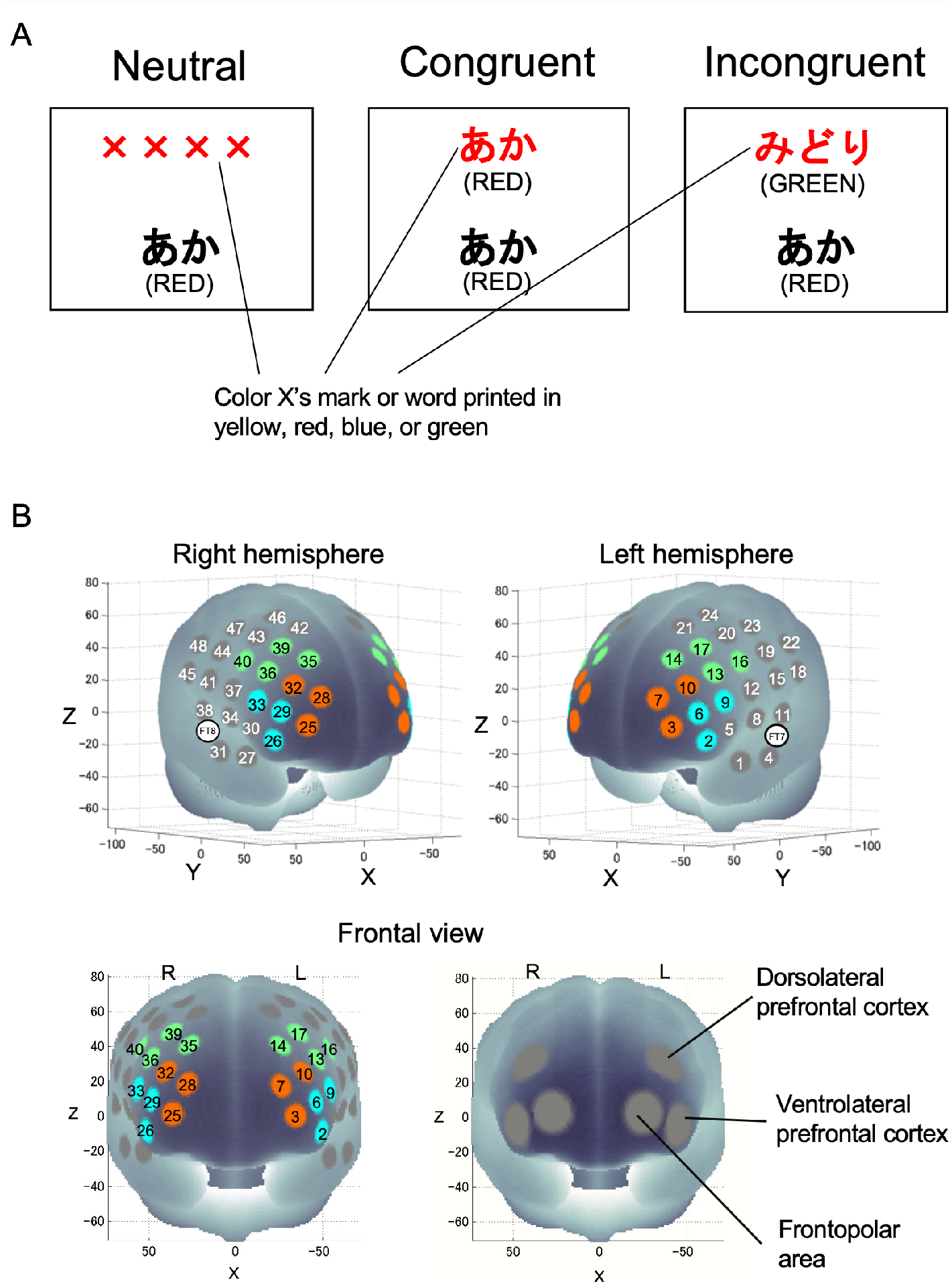
(A) Examples of the color-word Stroop test neutral, congruent, and incongruent conditions. The presented words were written in Japanese. English translations are shown in parentheses. (B) Spatial profiles of fNIRS channels used. Two sets of probe holders were placed to cover both lateral prefrontal activation foci as in our previous studies. Colors indicate each region of interest (ROI) in the lateral part of the PFC.

### fNIRS measurements

Cortical hemodynamic changes were monitored using the multichannel fNIRS optical topography system ETG-7000 (Hitachi Medical Corporation, Japan) with two wavelengths of near-infrared light (785 and 830 nm). The optical data from fNIRS was analyzed based on the modified Beer-Lambert Law as previously described. With this method, we were able to calculate signals reflecting the oxygenated hemoglobin (oxy-Hb) concentration changes in millimolar-millimeters (mM.mm). The sampling rate was set at 10 Hz. For fNIRS probe placement, two sets of 4 × 4 multichannel probe holders, which consist of 8 illuminative and 8 detective probes arranged alternately at an inter-probe distance of 3 cm resulting in 24 channels (CH) per set, were placed to cover both lateral prefrontal activation foci as in previous studies. The left probe holder was placed such that probe 5 (between CH4 and CH11) was placed over FT7, with the medial edge of the probe column parallel to the medial line. The right probe holder was symmetrically placed on the left hemisphere (Fig. 2B). We adopted a virtual registration method to register fNIRS data to Montreal Neurological Institute (MNI) standard brain space. With this method, we were able to place a virtual probe holder on the scalp by stimulating the holder’s deformation and by registering probes and channels onto a reference brain in the magnetic resonance image (MRI) database. We probabilistically estimated the MNI coordinate values for fNIRS in order to obtain the most likely microanatomical predictions for locations of the given channels as well as the spatial variability of the estimation. Finally, a MATLAB function was used to label the estimated locations in a macro-anatomical brain atlas.

### Analysis of fNIRS data

Prefrontal Oxy-Hb changes that occurred while performing the CWST were calculated as shown in our previous studies. Individual timeline data for each channel were preprocessed with a band-pass filter using a high-pass filter (0.04 Hz) to remove baseline drift and a low-pass filter (0.30 Hz) to screen out heartbeat pulsations. The motion artifacts were checked by using the optical topography analysis tools (POTATo) (Hitachi, Ltd., Japan). Then, channel-wise and subject-wise contrasts were calculated by the inter-trial mean of differences between peak (4-11 s after trial onset) and baseline (0-2 s before trial onset) periods (Byun *et al*., 2014*b*; Damrongthai *et al*., 2021; Kuwamizu *et al*., 2021). The contrasts obtained were subsequently subjected to a second level of random effects group analysis. This study adopted LBPA40, a widely used method among anatomical labeling systems, to combine 3 to 4 neighboring channels to form each region of interest (ROI). The regions included the left dorsolateral PFC (l-DLPFC; CHs 13, 14, 16, and 17), the right DLPFC (r-DLPFC; CHs 35, 36, 39, and 40), the left ventrolateral PFC (r-VLPFC; CHs 2, 6, and 9), the right VLPFC (r-VLPFC; CHs 26, 29, and 33), the left frontopolar area (l-FPA; CHs 3, 7, and 10), and the right FPA (r-FPA; CHs 25, 28, and 32). LBPA40 is considered valid because optical properties of neighboring channels are known to be similar. However, with this method, optical properties in different ROIs can cause systematic bias during statistical analysis. Therefore, we limited the analyses to ROI-wise and used a false discovery rate (FDR) to control the low proportion of false positives.

### Head acceleration measurements

An AS-20GB acceleration sensor was connected to a strain amplifier DPM-600A (both are products of Kyowa Electronic Instruments Co., Ltd.). This output signal was recorded by a personal computer after real-time conversion to a digital signal using PowerLab/16SP (ADInstruments). The sensor was attached on the top of the head with a lightweight bracket so that the acceleration detection direction was perpendicular to the floor. Very slow running was executed on a treadmill for two minutes. Vertical HA was measured and recorded while very slow running was executed on a treadmill for two minutes. The head moves up and down vertically once with each step, causing acceleration of the head. The incremental difference between the minimum and maximum acceleration values for ten steps were standardized (see Supplemental Material 3). HA was assessed separately after the SRUN condition because the accelerometer, which was rigidly attached to the head to maintain high validity, could interfere with the pleasant sensations of running or resting for 10 minutes.

### Statistical analysis

Psychological mood state was subjected to repeated-measures two-way analysis of variance (ANOVA) with session (CON, SRUN) and time (pre, post) as within-subject factors. Behavioral Stroop performance was first tested to determine whether there was significant Stroop interference using repeated-measures three-way ANOVA with condition (neutral, incongruent), session (CON, SRUN), and time (pre, post) as within-subject factors. Then, the effect of running on Stroop task performance was analyzed using repeated-measures two-way ANOVA with session (CON, SRUN) and time (pre, post) as within-subject factors. Cortical activation during pre-sessions for CON and SRUN, which were free from any effect of running, were first examined to determine whether the Stroop interference effect could be observed using one-sample *t*-test with FDR correction. Only significant ROIs for Stroop interference were subsequently analyzed for the effect of running on prefrontal activation using repeated-measures two-way ANOVA with session (CON, SRUN) and time (pre, post) as within-subject factors followed by FDR correction. Additionally, Pearson’s correlation coefficient was adopted to analyze the relationship between HA during SRUN and changes in variables to determine whether significant changes could be observed: [(post-session) - (pre-session)] for SRUN and [(post-session) - (pre-session)] for CON. The statistical significance level was set *a priori* at *p* < 0.05. The SPSS Statistical Package version 24 (SPSS, Inc., USA) was used for statistical analyses.

## RESULTS

### Verification of very slow running

To verify whether the participants could perform very slow running or a very light-intensity exercise, we monitored HR and RPE during exercise. During 10 min of very slow running, average HR and RPE were 104.84 + 7.61 bpm and 8.50 + 1.95 points, respectively, for males and 107 + 9.39 bpm and 8.68 + 1.27 points, respectively, for females. Based on the guideline of the American College of Sports Medicine, these values were determined to fall within the range of very light-intensity exercise. In addition, average treadmill speeds for males and females were 5.46 + 1.77 km/h and 3.86 + 0.87 km/h, respectively.

### Behavioral results

RT and ER had significant main effects of condition (neutral/incongruent; *F*(1, 23) = 63.59, *p* < 0.001 (Fig. 3A) and *F*(1, 23) = 20.03, *p* < 0.001 (Fig. 3B), respectively; repeated-measures three-way ANOVA), showing that the Stroop interference effect was basically found between the neutral and incongruent conditions in all sessions of this study. Next, the effects of running on Stroop interference RT and ER were examined. Stroop interference RT revealed significant interaction between session (CON, RUN) and time (pre, post) (*F*(1, 23) = 7.66, *p* < 0.05; repeated-measures two-way ANOVA; Fig. 3C), whereas Stroop interference ER did not reveal significant interaction (*F*(1, 23) = 2.09, *p* = 0.162). Finally, changes in Stroop interference RT were investigated and the results showed that Stroop interference RT after the SRUN session was significantly more reduced than that after the CON session (*t*(23) = 2.77, *p* < 0.05, Cohen’s *d* = 0.57; paired *t*-test; Fig. 3D).

**Figure 3.**
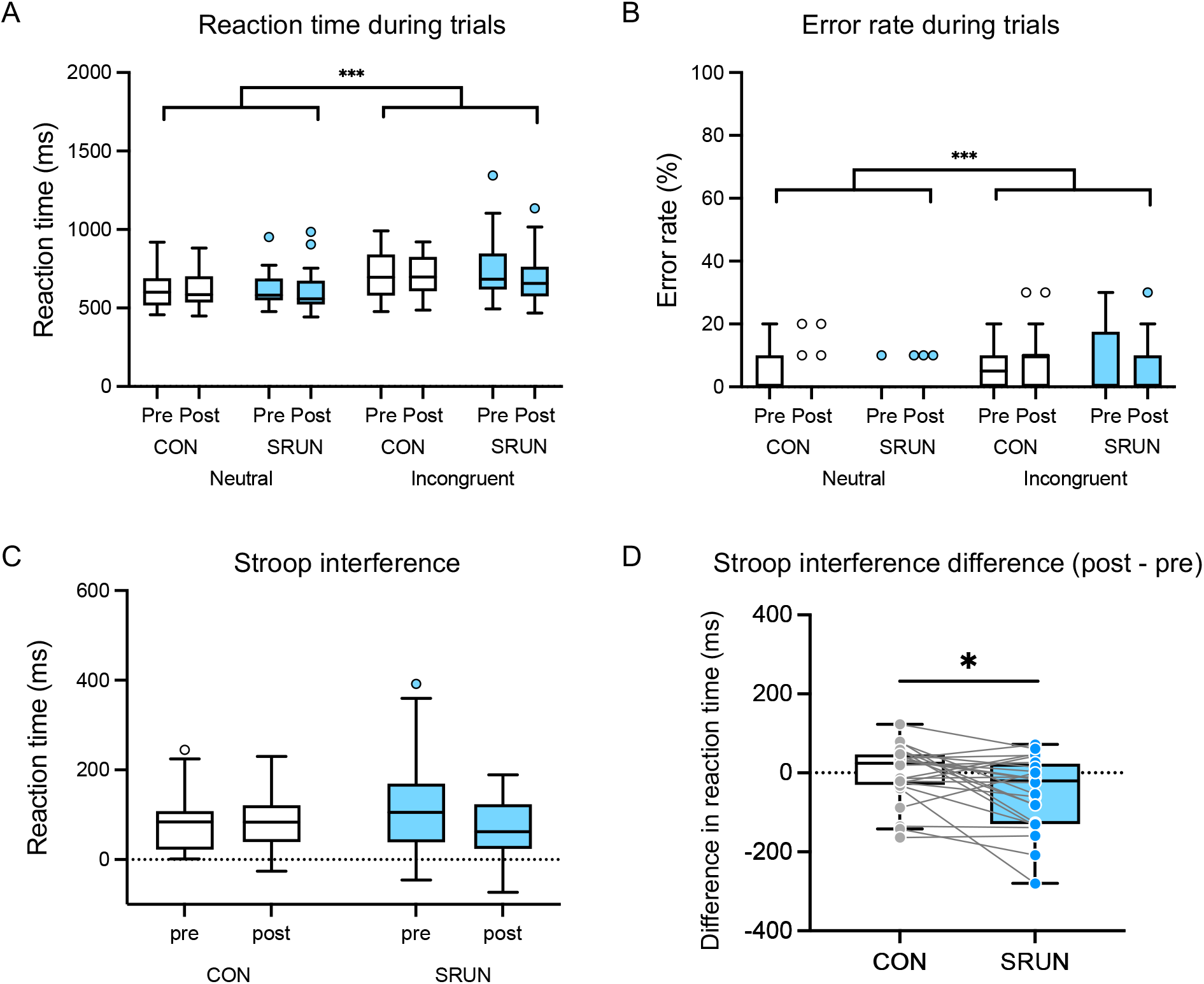
Comparison of Stroop task performance for (A) reaction time and (B) error rate between neutral and incongruent conditions. (C) Difference in Stroop interference [incongruent - neutral] between control and running sessions. (D) Contrast in Stroop interference difference ([incongruent - neutral of post-session] - [incongruent - neutral of pre-session]) between control and running. The box-and-whisker plots are drawn in the Tukey manner. Line plots represent individual data. *** = *p* < 0.001, * = *p* < 0.05

### Prefrontal activation

There were no significant differences in pre-session Stroop-interference-related cortical activation between CON and SRUN conditions in any ROIs. The pre-session data for the CON and SRUN sessions, which were free from any effect of exercise, were averaged and served as substrates for a ROI-wise analysis. Significant Stroop-interference-related cortical activations were found in 4 ROIs, l-DLPFC, l-VLPFC, l-FPA, and r-FPA (*p* < 0.05, one-sample *t*-test, FDR correction), indicating that the Stroop interference effect was observable in this study (Fig. 4).

**Figure 4.**
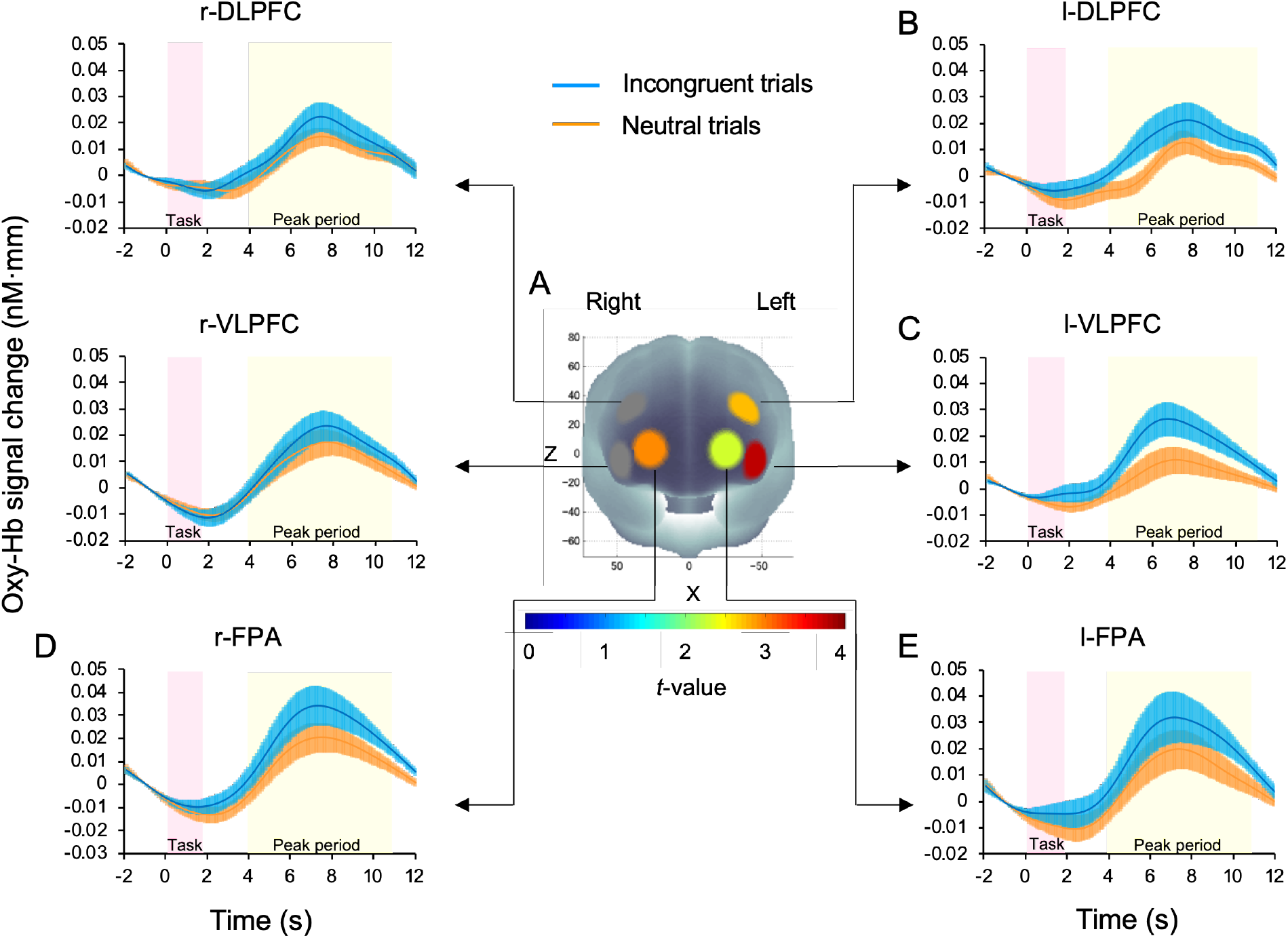
Cortical activation patterns during performance of the CWST for the CON and SRUN pre-sessions. Presented data are comparisons of averaged values between the CON pre-session and the SRUN pre-session. Baseline (2s before trial onset) was set at zero and peak periods were from 4 s to 11 s after trial onset. (A) *t*-map of (Oxy-Hb) signal change; *t*-values are as indicated by the color bar. Significant Stroop-interference-related cortical activations [incongruent - neutral] were found in 4 ROIs: the left dorsolateral prefrontal cortex (B), the left ventrolateral prefrontal cortex (C), the right frontopolar area (D), and the left frontopolar area (E). Data are mean + SE.

The effect of very slow running on prefrontal activation in these 4 ROIs was subsequently determined and the results reveal that Stroop-interference-related cortical activations had significant interactions between session (CON, SRUN) and time (pre, post) in the l-DLPFC (*F*(1, 23) = 7.73, *p* < 0.05) and the l-FPA (*F*(1, 23) = 7.34, *p* < 0.05) (repeated-measures two-way ANOVA). Finally, oxy-Hb change with Stroop interference in the l-DLPFC and the l-FPA were examined. The results reveal that the SRUN session had a significantly greater increase of oxy-Hb with Stroop interference than did the CON session in the l-DLPFC (*t*(23) = 2.78, *p* < 0.05, Cohen’s *d* = 0.57) and l-FPA (*t*(23) = 2.71, *p* < 0.05, Cohen’s *d* = 0.55) (paired *t*-test, FDR correction; Fig. 5).

**Figure 5.**
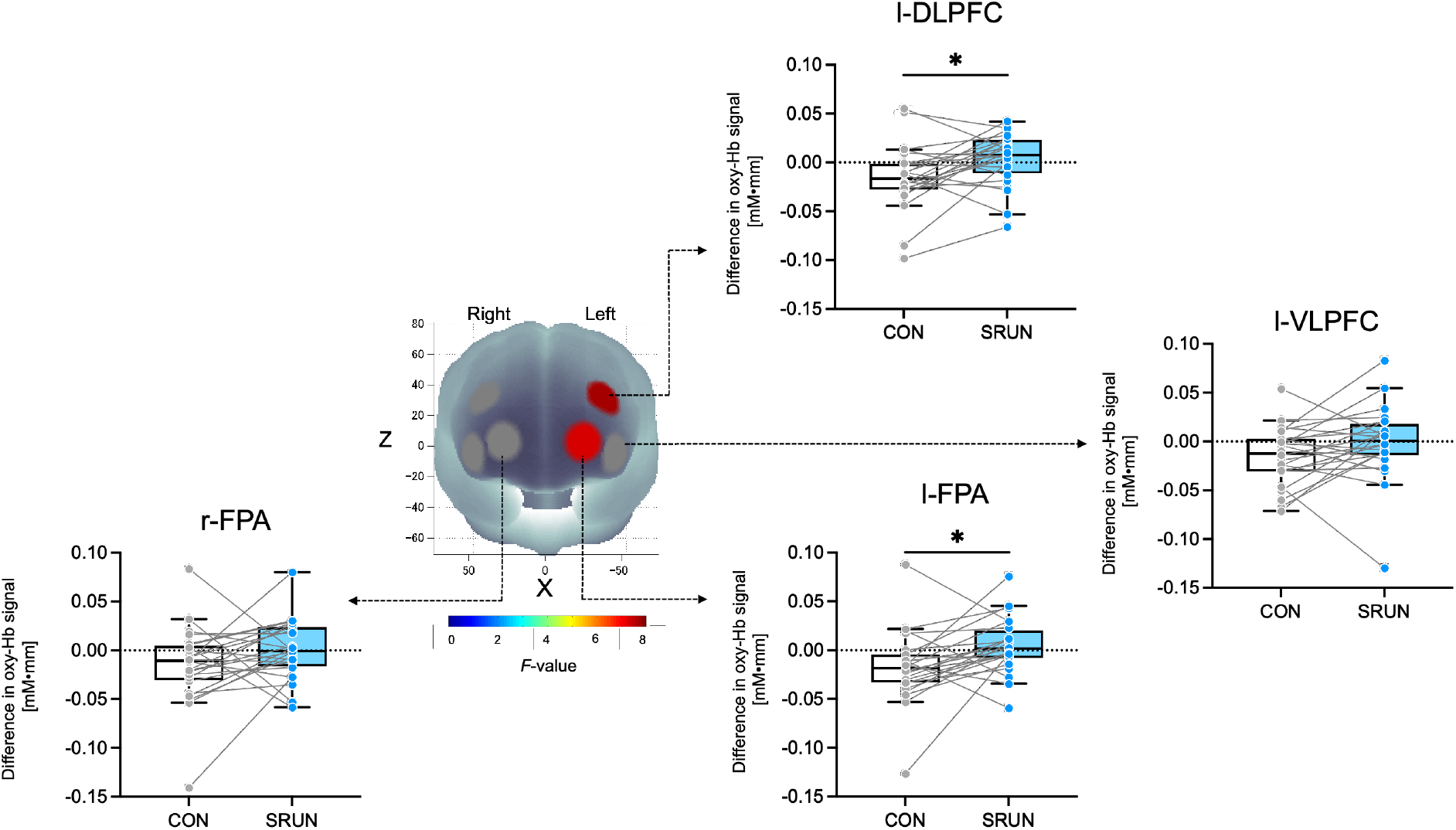
Very slow running elicits prefrontal activation in response to Stroop interference ([incongruent - neutral of post-session] - [incongruent - neutral of pre-session]). For the *F*-map of (Oxy-Hb) signal change, *F*-values are as indicated by the color bar. Among the 4 ROIs, significant differences are found in the left dorsolateral prefrontal cortex and the left frontopolar area. The box-and-whisker plots are drawn in the Tukey manner. Line plots represent individual data. \**p* < 0.05

### Psychological mood

Arousal and pleasure were found to have significant interactions between session (CON, SRUN) and time (pre, post) (*F*(1, 23) = 55.84, *p* < 0.001 and *F*(1, 23) = 22.96, *p* < 0.001, respectively; repeated-measures two-way ANOVA). Changes of arousal and pleasure were subsequently examined revealing that the SRUN session had a significantly greater increase of arousal and pleasure than did the CON session (*t*(23) = 7.47, *p* < 0.001, Cohen’s *d* = 1.53 [Fig. 6A] and *t*(23) = 4.79, *p* < 0.001, Cohen’s *d* = 0.98 [Fig. 6B], respectively; paired *t*-test).

**Figure 6.**
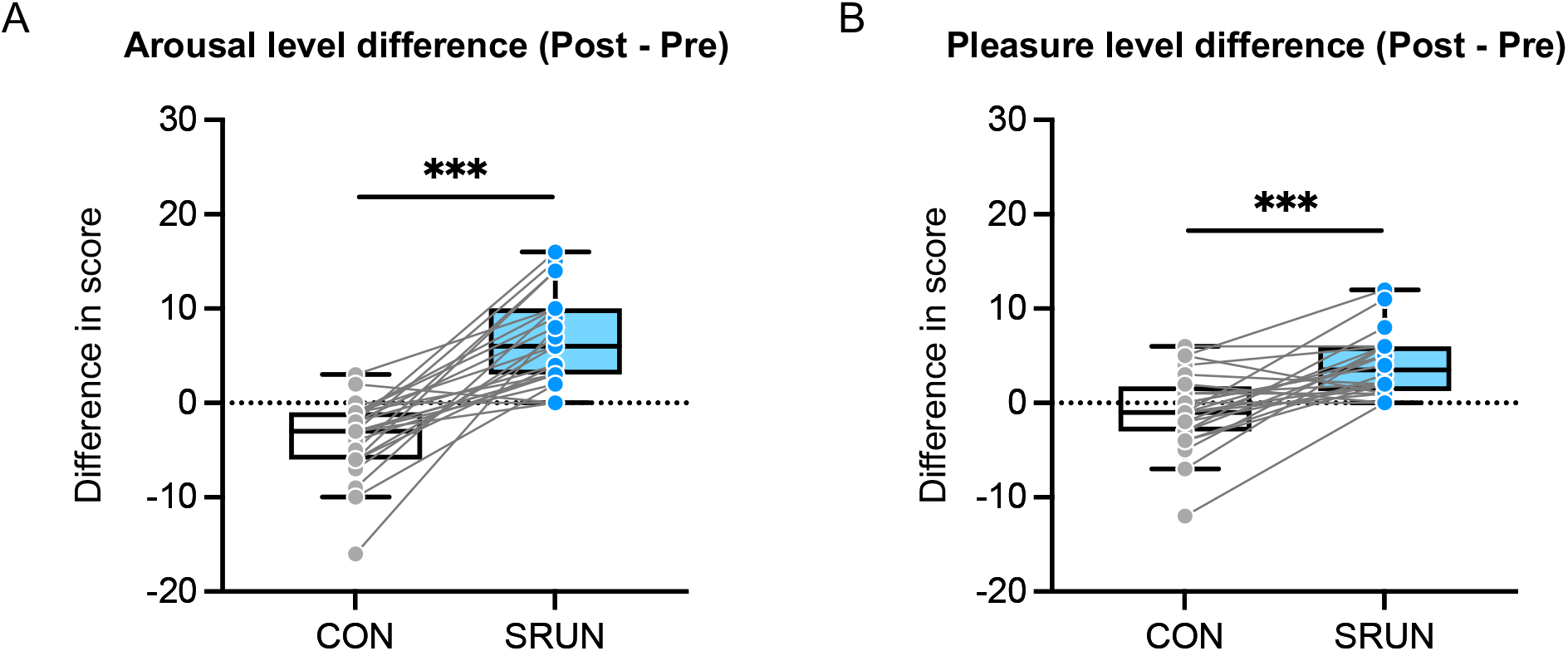
Comparison of mood change [(post-session) - (pre-session)] between CON and SRUN: (A) arousal level differences and (B) pleasure level differences. CON = control session, SRUN = very slow running session. The box-and-whisker plot is drawn in the Tukey manner. Line plots represent individual data. *** = *p* < 0.001

### Relationship between head acceleration and significantly changed variables with running

At the individual very slow running speed, the vertical HA averages in males and females were 2.14 + 0.48 G and 2.22 + 0.67 G respectively. HA influenced by very slow running was positively correlated with exercise-induced pleasure levels (*r* = 0.409, *p* < 0.05, Pearson correlation; Fig. 7). However, there was no significant association between HA and other variable changes induced by very slow running: Stroop interference reaction time (*r* = -0.03), l-DLPFC activation (*r* = -0.04), l-FPA activation (*r* = -0.34), and arousal level (*r* = -0.04); *p*-values for all variables were > 0.05, Pearson correlation. In addition, there was no significant correlation between HA and other potential covariate factors: treadmill speed (*r* = 0.20), 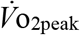 (*r* = 0.17), height (*r* = -0.05) and weight (*r* = -0.22): *p*-values for all variables was > 0.3, Pearson correlation.

**Figure 7.**
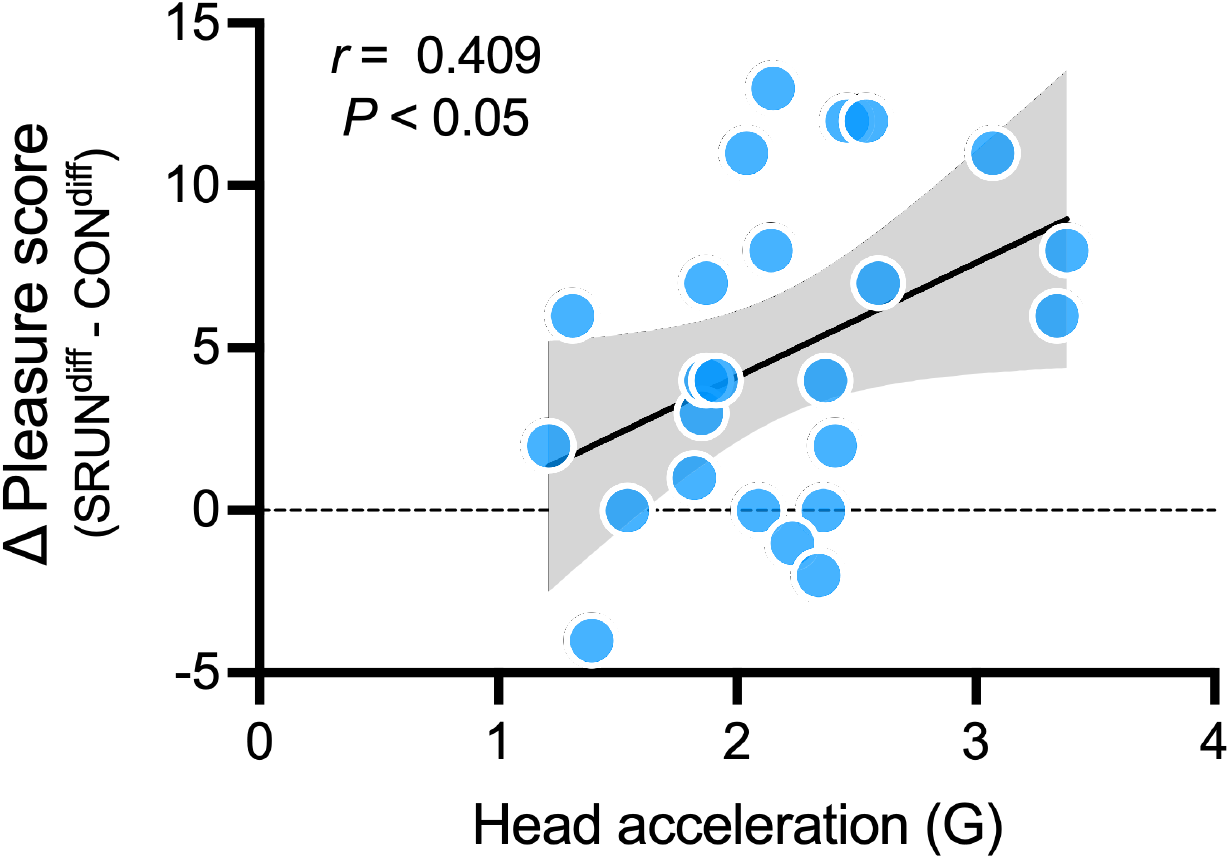
Correlation between running-generated head acceleration and running-enhanced pleasure level. The black line represents linear regression and the gray band represents 95% confidence bands.

## DISCUSSION

The current study confirmed the hypothesis that a single bout of very slow running, a very light-intensity exercise at 35% 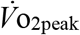, enhances executive function with activation in lateral prefrontal subregions. Moreover, very slow running elicited an improvement in mood and 17% of the variance in mood enhancement was explained by inter-individual HA. The current results suggest that acute very slow running has a positive effect on both executive function enhancement and mood improvement with minimal effort; part of this mechanism may originate from up-and-down oscillations of the body in addition to the comparable mechanisms of pedaling exercise.

Stroop interference (longer RT with higher ER in the incongruent condition compared to the neutral condition) was found across all sessions. Subsequently, the Stroop interference time was compared between conditions, and the results support the hypothesis that very slow running shortens Stroop interference time compared to resting control. This result is consistent with previous results on very light- or moderate-intensity pedaling (Yanagisawa *et al*., 2010; Byun *et al*., 2014*b*; Kujach *et al*., 2018) and moderate-intensity running (Damrongthai *et al*., 2021). Therefore, the present findings build upon the results of previous studies by suggesting that even very slow running benefits executive function.

Subsequently, we focused on investigating the neural substrates for Stroop interference. In response to Stroop interference (Incongruent - Neutral), Oxy-Hb signal significantly increased in the 4 ROIs: l-DLPFC, l-VLPFC, l-FPA, and r-FPA during pre-sessions. These results are consistent with previous neuroimaging studies, including fNIRS, electrostimulation, and lesions, which have reported that Stroop interference is consistently associated with the PFC, and more specifically l-PFC (Perret, 1974; MacDonald *et al*., 2000; Yanagisawa *et al*., 2010; Byun *et al*., 2014*b*; Frings *et al*., 2018; Kuwamizu *et al*., 2021). The effect of running on prefrontal activation was subsequently determined, and the results show that very slow running elicits significant increases in activation in the l-DLPFC and l-FPA. The l-DLPFC, a brain locus implicated in inhibitory control and mood regulation, had increased Oxy-Hb response to Stroop interference (Leung *et al*., 2000; MacDonald *et al*., 2000; Song & Hakoda, 2015). Furthermore, the FPA, a brain region which generally activates with the DLPFC in response to tasks involving manipulation and monitoring, such as planning for action, has been found to have significant activation on the left side (Christoff & Gabrieli, 2000). These results are consistent with a previous study, which demonstrated that 10 min of very-light-intensity pedaling elicits activation of the l-DLPFC and l-FPA with Stroop interference processing (Byun *et al*., 2014*b*; Kuwamizu *et al*., 2023). It is suggested that the neural basis on which very slow running improves executive function is similar to that of pedaling exercises at the same intensity. On the other hand, previous studies on acute moderate-intensity running have shown extensive activation of the r-PFC in addition to the left side with Stroop interference processing (Damrongthai *et al*., 2021). This difference in prefrontal activation patterns may depend on exercise intensity (very slow vs. moderate). Very slow running may have minimal hypothalamus–pituitary–adrenal axis stress responses and distinct differences in the level of activation of the arousal system compared to moderate or more intense efforts (Ohiwa *et al*., 2006; Soya *et al*., 2007), which may be related to the prefrontal activation area elicited by running. This requires further investigation.

Our previous study showed that moderate-intensity running causes mood changes related to arousal and pleasure that coincide with the facilitation of PFC function (Damrongthai *et al*., 2021). Based on these previous findings, we tested whether acute very slow running would improve mood. As we hypothesized, very slow running improved both arousal and pleasure levels compared to resting control as well as the moderate-intensity running of the previous study did (Damrongthai *et al*., 2021). Previous pedaling studies have shown that very light-intensity exercise increases psychological arousal (Byun *et al*., 2014*b*; Suwabe *et al*., 2018), possibly associated with the catecholaminergic system which originates from the locus coeruleus (Kuwamizu *et al*., 2022, 2023; Yamazaki *et al*., 2023). The current study also attempted to measure pupillometry and confirmed pupil dilation during running. However, the data was limited to a small sample size due to device errors related to body movement (see Supplemental Material 4). Understandably, the brain’s catecholaminergic arousal system is activated during very slow running, comparably to pedaling, and might contribute to the elicitation of prefrontal cortex activation and to improved executive function.

Interestingly, previous studies have shown that very light-intensity pedaling exercise increases arousal but has only a small effect on pleasure levels. Our exploratory analysis suggested that the positive impact of very slow running on pleasure exceeded the impact produced with a comparable intensity pedaling (Supplemental Material 5). This increase in pleasure level seen with very slow running may be characteristic of this exercise. In addition, HA, a property of running, during very slow running, was examined. Interestingly, the magnitude of HA was positively correlated only with changes in pleasure levels induced by running, and this may indicate a running-specific effect, as discussed above. Additional experiments confirmed that HA does not vary much during pedaling exercise, even if participants pedal at a high rate (Supplemental Material 6). This suggests that, in addition to the common exercise effects of pedaling and running (oxygen uptake level, heart rate, arousal level, and RPE are common to the two exercises), rhythmic up-and-down oscillations contribute to mood improvement. This is a new and challenging hypothesis, and the exact neural circuitry involved in the positive effect of HA on mood has not been identified and needs further investigation. A recent animal study demonstrated that HA modulates behavior and the serotonergic system in the prefrontal cortex (Ryu *et al*., 2020). As the brain’s serotonergic system is involved in emotion regulation, this is one candidate mechanism of running- and head-acceleration-induced mood enhancement. This mechanism may work in addition to the arousal system alone (i.e., with no HA) and may form a running-specific arousal state. As other candidate factors, physiological changes due to foot strike (Palatini *et al*., 1989; Lyngeraa *et al*., 2013) and high tempo rhythm sensation with foot beats might also be related to mood changes and PFC activity (Fukuie *et al*., 2022). The current results represent a starting point, and further research is expected to determine how HA during running correlates with positive mood changes. It would be interesting to investigate the effects of dose-response for head acceleration and other forms of rhythmic up-and-down oscillations such as jump rope (Yamashita & Yamamoto, 2021).

While head acceleration may play a role in mood alterations, it does not correlate with the improvements in executive function and activation of the lateral prefrontal subregions induced by running. Interestingly, even with minimal head acceleration during pedaling, it has been shown to positively enhance cognitive aspects, especially prefrontal executive function and hippocampal memory (Byun *et al*., 2014*b*; Suwabe *et al*., 2018). The lack of significant findings linking head acceleration to enhanced prefrontal executive function implies that it might not be a critical factor in the cognitive benefits derived from exercise benefits.

This study aimed to determine the lowest intensity/speed of running that would have a positive effect on prefrontal function and mood. The participants’ running speeds ranged from about 3 to 6 km/h. This speed is below the transition from walking to running (around 7 km/h) and is the lowest speed at which the form of running can be maintained. However, this study is the first to show that running, even at its lowest speed, is sufficient to induce a positive mood and to promote prefrontal executive function. Although this study included only healthy young adults, future validation with other populations will increase its generalizability. Even very light-intensity pedaling exercise, when maintained consistently, improves PFC function in older adults (Byun *et al*., 2023). It will be an interesting challenge to see if a long-term intervention of very slow running outperforms pedaling exercise.

In conclusion, the current study successfully determined the beneficial effect of a 10-min single bout of very slow running on improving mood and executive function with lateral prefrontal neural activation. This discovery of the positive relationship between head acceleration and improved mood may be one of the underlying mechanisms through which very slow running promotes mental health. To this end, these findings contribute to societal wellbeing by encouraging people with various health conditions to keep physically and mentally active, while also maintaining their own safety, through very slow running.

## ADDITIONAL INFORMATION

### Supplemental Material

Supplemental materials are available on the https://osf.io/wb49v/.

### Data availability statement

All data that supports the findings of this study are available from the corresponding author by request with no restrictions.

### Author contributions

C.D., R.K., K.A., and H.S. contributed to the design of the study. C.D., R.K., Y.Y., N.A., D.L., and K.A played a role in collecting data. Data analysis was done by C.D., R.K., Y.Y., K.A., and H.S. The manuscript was written by R.K. and C.D., edited by Y.Y., K.A., K.B., F.T., W.C., M.A.Y. and H.S. H.S. contributed to forming the concept of the study and funding acquisition. All authors have read and approved the final version of the manuscript.

## Acknowledgements

The authors thank the members of the Laboratory of Exercise Biochemistry and Neuroendocrinology, University of Tsukuba for their assistance in data collection. Also, they express their gratitude to M. Noguchi (ELCS English Language Consultation, Japan) for helping with the manuscript.

